# Diversification of Disease Resistance Receptors by Integrated Domain Fusions in Wheat and its Progenitors

**DOI:** 10.1101/695148

**Authors:** Ethan J. Andersen, Madhav P. Nepal

## Abstract

Pathogenic effectors inhibit plant resistance responses by interfering with intracellular signaling mechanisms. Plant Nucleotide-binding, Leucine-rich repeat Receptors (NLRs) have evolved highly variable effector-recognition sites to detect these effectors. While many NLRs utilize variable Leucine-Rich Repeats (LRRs) to bind to effectors, some have gained Integrated Domains (IDs) necessary for receptor activation or downstream signaling. While a few studies have identified IDs within NLRs, the homology and regulation of these genes have yet to be elucidated. We identified a diverse set of wheat NLR-ID fusion proteins as candidates for NLR functional diversification through ID effector recognition or signal transduction. NLR-ID diversity corresponds directly with the various signaling components essential to defense responses, expanding the potential functions for immune receptors and removing the need for intermediate signaling factors that are often targeted by effectors. ID homologs (>80% similarity) in other grasses indicate that these domains originated as functional, non-NLR-encoding genes and were incorporated into NLR-encoding genes through duplication. Multiple NLR-ID genes encode experimentally verified alternative transcripts that include or exclude IDs. This indicates that plants employ alternative splicing to regulate IDs, possibly using them as baits, decoys, and functional signaling components. Future studies should aim to elucidate differential expression of NLR-ID alternative transcripts.

## 1. INTRODUCTION

Plant innate immune systems utilize specialized receptor proteins to detect pathogens [1]. Nucleotide-binding, Leucine-rich repeat Receptor (NLR) proteins detect pathogen effectors that would otherwise inhibit host resistance responses [2]. In order to detect the hundreds of pathogens and pests, immune receptors must be able to respond to many elicitors. To accomplish this, NLRs have radiated to form a diverse family of resistant genes in plants [3]. Much of this diversity is accomplished by gene duplication and variation in NLR Leucine-Rich Repeats (LRRs), which allow NLRs to bind to new effectors [4]. Diversification has led to the formation of networks of sensor and helper NLRs, with some NLRs dimerizing to initiate signaling [5–8]. As another form of diversification, some NLRs have gained extra domains that may assist the receptor in pathogen recognition or in resistance signaling. These domains, called Integrated Domains (IDs), resulted from a fusion of NLR and functional domains also involved in resistance, as outlined by the integrated decoy/sensor model [9,10]. ID diversity has been probed across many plant genomes, revealing a diversity of domains associated with potential roles in resistance [11].

The major objectives of this study are to identify wheat NLR-ID fusion proteins, infer their function, and assess their homology in wheat relatives and other selected monocot species. We also propose a method plants utilize for NLR-ID regulation, which became apparent while manually assessing the transcript and protein sequences. Understanding the evolution of NLR-ID fusions provides a unique perspective on NLR diversification, which often gets ignored by large-scale analyses of NLR gene family evolution. These findings will improve our understanding of how NLRs diversify to oppose various pathogenic molecular weapons.

## 2. RESULTS

### 2.1. IDs in Wheat

We have identified wheat NLR proteins with IDs that potentially function as molecular baits, decoys, or signal transduction factors. Wheat NLRs possess a diverse set of IDs, the most commonly occurring are kinase and DNA-binding domains. Figure 1 shows the average location of these domains relative to protein length, averages calculated from every NLR-ID occurrence of that domain. Kinase domains are generally located in the N-terminal half of the protein with tyrosine kinase domains generally in the middle of the protein sequence. The more diverse class of DNA-binding domains vary by domain type, present in both the N- as well as C-termini. For example, Myb-like and BED zinc finger domains are generally at the N-terminus, while B3 and WRKY domains are at the C-terminus. Many other IDs located at the C-terminus have potential roles in signaling, such as calmodulin-binding, jacalin-like lectin, thioredoxin, and ubiquitin-conjugating.

**Figure 1.**
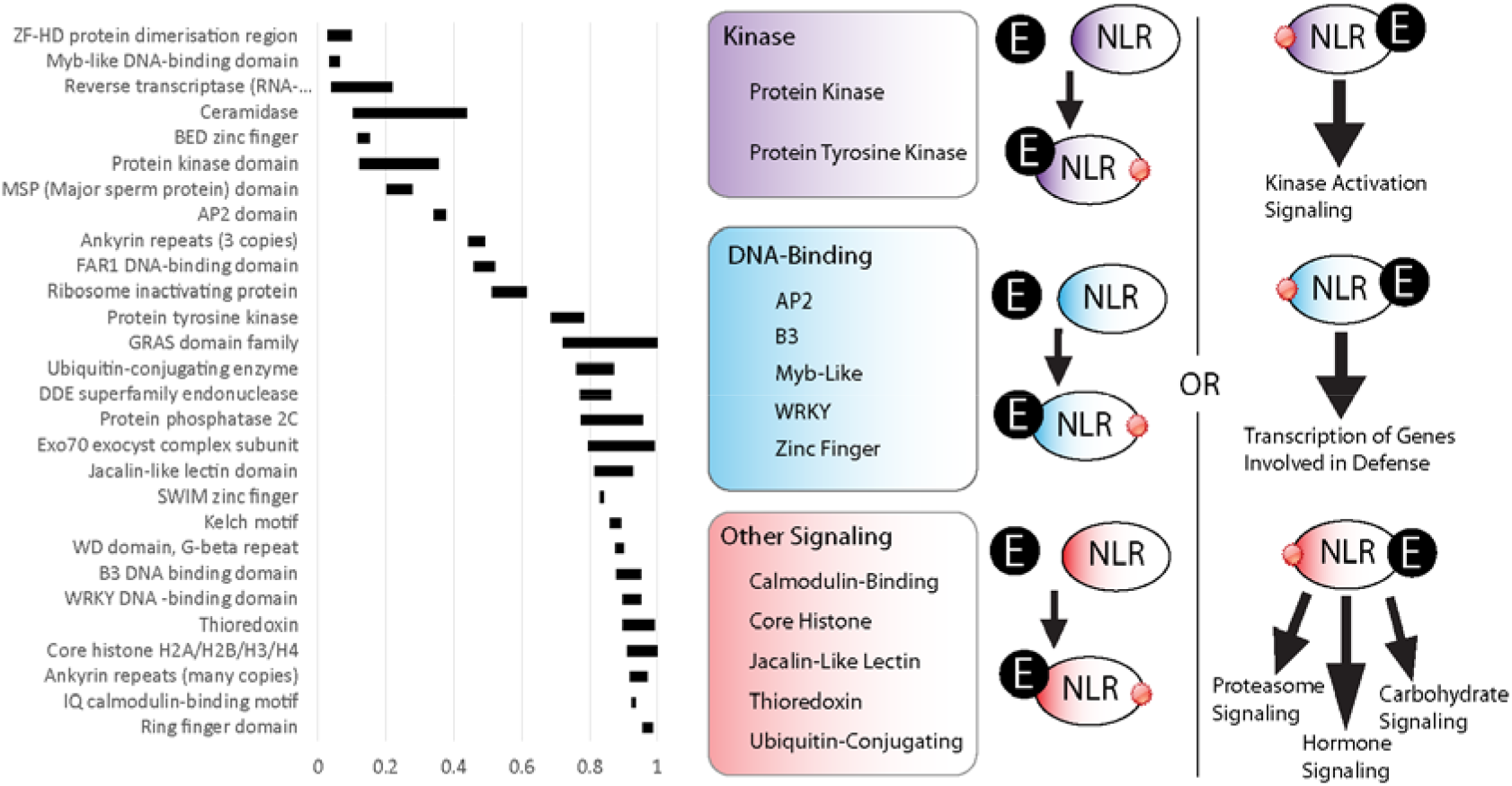
Integrated Domain (ID) locations, indicated by black rectangles, are shown within NLRs relative to protein length (0-1). IDs were grouped into functional categories, based on their potential involvement in kinase, DNA-binding, or other signaling activities (shown in purple, blue, and red, respectively). Schematic diagrams representing potential functions for these NLR-IDs are included with pathogenic effectors represented by black circles (labeled as ‘E’) and NLR-ID proteins as ovals color coded by ID type (i.e. ‘Kinase’, ‘DNA-Binding’, or ‘Other Signaling’). The diagram includes representations of both effector-bait interaction (left) and NLR-ID signaling (right) that these domains may be involved in. Red circles at the sides of NLRs indicate activated NLR proteins.

### 2.2. ID Homology

Many *Triticum aestivum* (TA) IDs share homology with proteins in distantly related monocots. Figure 2 shows the wheat accessions with high percent identity (above 70%), grouped by ID type and homolog species. The vast majority of these homologs do not contain NB-ARC domains, indicating a recent fusion. ID homologs in distant relatives generally were not NLR proteins. While other plants also possess NLR-ID fusions, many are lineage specific and are not conserved across diverse species. Barley, a close relative of wheat, possessed many of the same NLR-ID fusion proteins as in wheat, with 68.5% of ID homologs in barley also possessing NLRs. The two progenitors of wheat with sequenced genomes, *T. urartu* and *A. tauschii*, also possess wheat’s NLR-ID fusions. Of these progenitor homologs, 40.8% matched the subgenome expected based upon the known progenitor-subgenome relationships. Genomes investigated for homology include: *Aegilops tauschii* (AT), *Amborella trichopoda* (AmT), *Arabidopsis thaliana* (ArT), *Brachypodium distachyon* (BD), *Hordeum vulgare* (HV), *Musa acuminata* (MA), *Oryza sativa* (OS), *Setaria italica* (SI), *Triticum urartu* (TU), and *Zea mays* (ZM). Genomic data was not available for *Aegilops speltoides* (AS).

**Figure 2.**
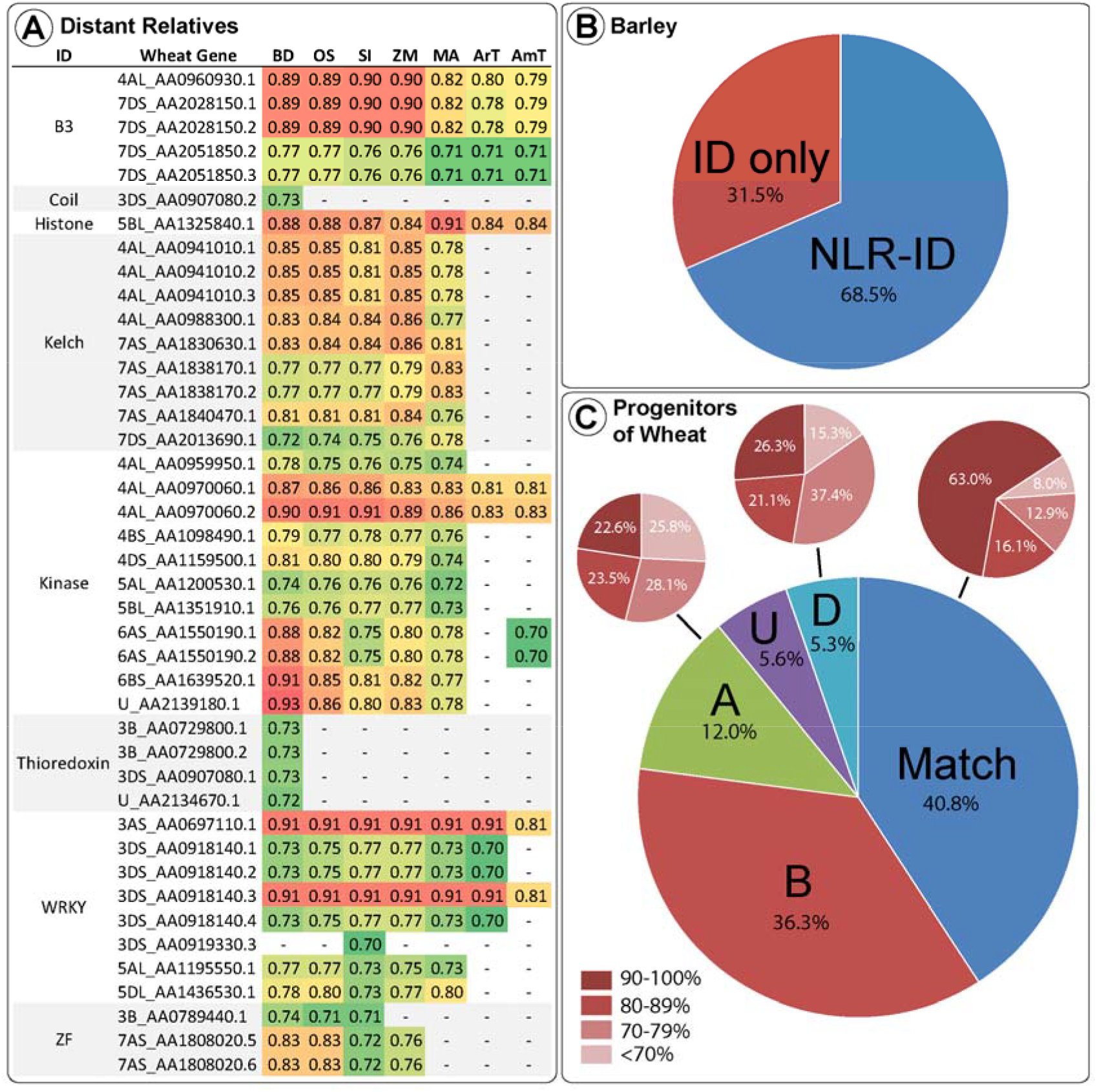
Wheat IDs and their homologs in wheat progenitors and other divergent monocot species are shown, including *Arabdopsis* (model plant species) and *Amborella* (basal known Angiosperm). **(A)** Sequence similarities above 70% are shown between wheat IDs and their homologs in Brachypodium (BD), rice (OS), foxtail millet (SI), maize (ZM), banana (MA), *Arabidopsis* (Art), and *Amborella* (AmT). Wheat accession names are shortened to only include the chromosome arm and the last digits unique to each transcript. (**B**) Barley ID homologs possessing and lacking NLR domains are compared. (**C**) Mapping of homologs among wheat and wheat progenitors is displayed – a match between the progenitor and subgenome (labeled ‘Match’); subgenome A protein was more similar to an *Aegilops tauschii* sequence (labeled ‘A’); subgenome D protein was more similar to a TU sequence (labeled ‘D’); sequence was from the B subgenome with the unavailable AS progenitor (labeled ‘B’), or the accession subgenome is unknown (labeled ‘U’). Also, the level of homology between the pairs is demonstrated for ‘Match’, ‘A’, and ‘D’, with dark red corresponding to the proportion of sequences with high similarity (>90%) and lighter red corresponding to lower similarity (<70%).

### 2.3. NLR-ID Regulation

Some NLR-ID genes encode alternative transcripts that omit IDs or other domains, such as transmembrane helices. Figure 3 illustrates a consolidation of all wheat NLR-IDs in which another transcript of the same gene excluded the ID. Alternative splicing of this kind would allow plants to regulate the use of IDs by including or excluding exons containing them. Similar characteristics were also observed in barley transcripts, indicating a conserved use of alternative splicing. Alternative transcripts may also be found in wheat progenitors, which currently lack available data. Expression data from the Wheat Gene Expression Atlas and NCBI shows differential expression between these alternative transcripts, which are shown in Figure S1 and **Table S1**.

**Figure 3.**
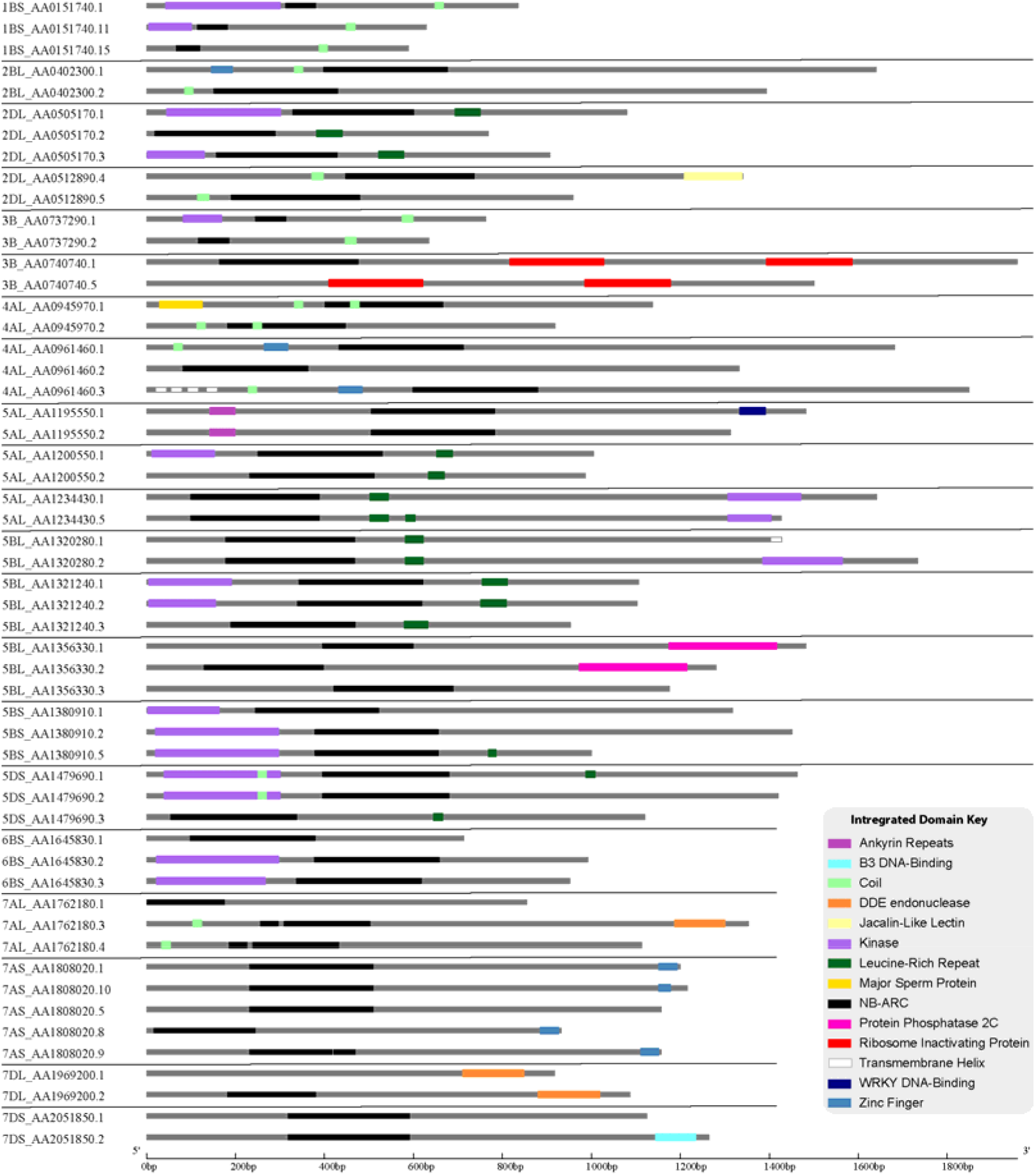
Wheat NLR-ID genes that encode alternative transcripts excluding or truncating IDs or other NLR domains are shown. Grey bars span the exon length in base pairs, black bars represent NB-ARC domains. Color segments annotate ID locations by domain type, which are defined in the integrated domain key. The scale, in base pairs, is shown along the bottom of the figure and black bars separate each set of transcripts. Wheat accession names are shortened to only show the chromosome arm, last code of gene name, and transcript number.

## 3. DISCUSSION

### 3.1. IDs Augment NLR Function Through Signaling and Recognition

Kinase and DNA-binding IDs likely function as signaling domains that help NLRs initiate defense responses. Current models of NLR function describe a conformational shift triggered when pathogenic effectors bind to the C-terminal LRR, causing the NB-ARC to exchange ADP for ATP, opening the protein up for the N-terminus to initiate further signaling [12–14]. LRRs, as highly variable domains of repeating Lxx amino acid residues, allow defense receptors to bind to diverse elicitors. The NB-ARC, as a P-loop-containing nucleoside triphosphate hydrolase, functions in hydrolysis of beta-gamma phosphate bonds in ATP, binding to phosphates using the Walker A (P-loop) motif and to magnesium ions necessary for catalysis by Walker B motifs [15]. This release of energy from ATP hydrolysis drives protein conformational change, allowing N-terminal domains (i.e. TIR or CC) to trigger downstream signaling. Kinase IDs we found in wheat NLRs could initiate signaling through phosphorylation of transcription factors or other kinases (i.e. MAPK). Sarris et al. (2016) also found an abundance of NLR-kinase fusions, which possibly retain their biochemical activity [11]. DNA-binding domains could move directly to the nucleus upon activation, binding to promoters of pathogenesis-related (PR) genes to recruit transcription machinery. IDs that likely bind to DNA include: AP2, B3, Zinc Finger, Myb, and WRKY domains, which have been shown to play roles in pathogen resistance [16,17]. The *Arabidopsis* NLR gene AT4G12020 has been identified both as MAPKKK11 [18] and a TNL resistance gene [19], containing WRKY DNA-binding sites and a protein kinase domain. This gene is a homolog of SLH1, which has been associated with hypersensitive response, possibly guarding a pathogen effector target [20]. Many NLR-ID fusion proteins contain transmembrane (TM) domains or nuclear localization signals (NLSs). Several proteins have multiple TM domains, with proteins like 3B_AA0787000 containing seven, characteristic of other TM proteins. NLSs indicate that DNA-binding domains may functionally interact with DNA as transcription factors.

In addition to signaling, some IDs may play direct roles in effector recognition as effector-binding domains or bait domains that mimic effector targets. Jacalin-like lectin domains, for example, bind to carbohydrates and can recognize carbohydrates that originate directly from pathogens or from damage incurred during infection [21–23]. Mannose-binding lectin domains were also found in NLRs, associated with disease resistance [24], along with “Wall-associated receptor kinase galacturonan-binding” and “Cleavage site for pathogenic type III effector avirulence factor Avr” domains. Lectin domains may distinguish proteins as helper NLRs, with carbohydrates acting as signals to initiate NLR activation. Other domains may play roles in effector recognition as bait domains that resemble effector targets. The resistance protein RRS1 becomes activated when an integrated WRKY domain interacts with *Ralstonia solanacearum* effector PopP2 and *Pseudomonas syringae* pv. *pisi* effector AvrRps4, effectors that otherwise target WRKY transcription factors [25,26]. Wheat NLR-WRKY fusions share homology with WRKY16, WRKY19, WRKY46, and WRKY54/70, with potential roles as targets, especially WRKY46, which is associated with bacterial resistance [11]. Accession 5DL_AA1436530 contains two variants of WRKY domains (WRKY and WSKY), possibly providing diverse baits for effectors. Accession 2BL_AA0441310 encodes a protein with separate AP2 and BED zinc finger domains, either allowing it to bind to separate promoters or as bait for more than one effector. Some bait proteins, such as PBS1, are kinases that pathogen effectors target for degradation, increasing the utility of NLR-kinase fusions. The Rosetta stone theory describes this association between fusions and linkage between protein function [27]. Several proteins with IDs and NB-ARCs do not contain LRRs, which would not be required for activation since baits have replaced LRRs in function.

The activity of IDs as baits is further supported by ID diversity, which corresponds to the diversity of defense regulatory components. IDs found in NLRs are also found in proteins that effectors target to interfere with defense. Several domains correspond to proteins involved in resistance signaling: calmodulin-binding (calcium signaling), Gibberellic acid insensitive (GAI) Repressor of GAI And Scarecrow (GRAS; gibberellin signaling), and ethylene responsive element binding (ethylene signaling). Several different domains contain IDs associated with the proteasome or ubiquitin, also involved in regulating resistance: protease subunit, proteasome component signature, cullin-repeat, RING/U-box, ubiquitin conjugating enzyme, and WD domains. Some IDs contain domains associated with regulation of DNA expression: core histone and chromatin organization modifier. Other IDs correspond to proteins involved in resistance responses: ribosome inactivating and ricin domains (disrupt ribosome activity), thioredoxin and kelch (oxidase activity in reactive oxygen species production), alpha subunit of tryptophan synthesis (synthesis of antiherbivory and antimicrobial compounds), Exo70 exocyst complex subunit (transport of antimicrobial compounds out of the cell), and DDE endonuclease (apoptosis). A few other domains are likely associated with pathogen components: major sperm protein (nematode sperm function, targeted by plant RNA interference), FNIP (found in *Dictyostelium discoideum*), and reverse transcriptase (inhibition of viral infection). Additional viral IDs include: RNA-binding/recognition, retrovirus zinc finger-like domain, and integrase domains. IDs may also be associated with pathogen-derived resistance and RNAi that plants use to inhibit viruses and other pathogens. Other studies support this diversity in IDs, showing similar results in other plant species [11,28].

### 3.2. IDs Originate as Functional Domains and Close Relatives Share NLR-IDs

Bread wheat is an allohexaploid species resulting from two separate hybridization events and substantial artificial selection [29]. As a monocot, wheat shares distant relationships with other members of the family Poaceae (i.e. BD, ZM, SI, and OS). Naturally, much strong similarities exist between wheat and other members of the Triticeae tribe, including HV and wheat progenitors TU, AS, as well as AT. ID homologs in distant relatives generally do not contain NB-ARCs, indicating relatively recent origin of NLR-ID fusions. IDs with high percent similarity to homologs, indicative of functional retention, include: proteasome subunit, B3 DNA-binding, WRKY DNA-binding, core histone, protein kinase, and kelch motif. Other domains with moderate similarity include: jacalin-like lectins, ribosome inactivating protein, BED zinc finger, SWIM zinc finger, ZF-HD protein dimerization region, zinc knuckle, protein phosphatase 2C, tyrosine kinase, thioredoxin, major sperm protein, reverse transcriptase, and DDE endonuclease. Homologs may be obscured since mutations accumulate in regions not essential for function or effector-bait interaction, causing divergence from the original sequences and making homology difficult to assess. Some mutations may increase functionality of NLR-IDs, since the original ID sequence was functionally optimized within a different protein. Some IDs may serve as baits for multiple targets, if targets possess similar modification/cleavage sites (e.g. similar WRKY domains).

Many IDs showed high homology in distant relatives. Kinase domains of up to 300 amino acids in length were over 80% similar to homologs. DNA-binding domains also had high homology in distant relatives. WRKY DNA binding domains present in wheat and progenitors have 90.5% similarity to several non-NLR genes in AT, BD, MA, OS, SI, and ZM. Many other IDs in wheat and its progenitors share >80% similarity with homologs in SI, ZM, BD, OS, AT, MA, and AmT. These results concur with previous investigations into IDs, where conserved IDs were identified in diverse plant species [11,30]. Some wheat proteins are very similar to their homologs in TU and AT, whereas others provide examples of one species diversifying from the other two. The histone ID in wheat protein 5BL_AA1325840 (approximately 100 amino acids) shares strong homology (>80%) with proteins in MA, BD, OS, SI, ArT, AmT, and ZM, a recent fusion not present in wheat relatives. Greater than 90% similarity was observed between the 182 amino acid long F775_12304|EMT01588 proteasome subunit domain and proteins in BD, OS, SI, and ZM. While this indicates that these accessions are close homologs, none of the other accessions have NB-ARCs, only peptidase, proteasome subunit, and nucleophile aminohydrolase domains. HV, TU, and TA homologs to this domain, while matching the sequence 100%, do not have NB-ARCs, indicating a very recent duplication and then fusion, after the hybridization of hexaploid wheat. Kelch motif IDs were found in one TU, three AT, and six TA proteins. Interestingly, only one of the TA proteins is in the D subgenome and 5 in the A subgenome, when the opposite would be expected based upon subgenome origin.

While domain homology in distantly related species indicates functional origins of IDs, homologs identified in close relatives (i.e. HV) and wheat progenitors (TU and AT) indicate recent fusions and duplications. Unlike distant relatives, barley shares many NLR-ID fusions with wheat. This indicates that many of wheat’s NLR-IDs happened before the divergence of barley and wheat progenitors. Since this divergence, wheat and barley have independently gained and lost NLR-ID proteins. Many TU and AT proteins are almost identical to proteins encoded by TA genes within the A and D subgenomes. IDs within wheat’s B subgenome often originate from AS and do not have 100% homologs in TU and AT. In select cases, similarity was found between NLR-IDs and functional domains from non-NLR proteins, indicating potential NLR-ID fusions since the formation of wheat. Conversely, some close wheat relatives share homology with distantly related NLR-ID fusions, such as F775_00546|EMT17242 and Si008625m, with an 84.6% similarity (991 identical sites) between their whole sequences, both with NB-ARCs and kinase domains. Unexpectedly, many proteins were found in wheat but not a relative, or vice versa. These domains include: histone, ribosome inactivating protein, calcium signaling, cleavage for type III effectors, RNase H type, P450, antibiotic synthesis, and ubiquitin conjugating enzyme. Other domains were found in greater or lower numbers in wheat compared to its progenitors, such as DDE endonuclease and reverse transcriptase, indicating loss or duplication in one genome.

### 3.3. Plants Use Alternative Splicing to Regulate NLR-IDs

Many NLR-ID protein-encoding genes possess multiple transcripts, some of which lack IDs or truncate domains within protein. Examples found in wheat are displayed in Figure 3. This indicates that plants may use alternative splicing to regulate the use of IDs within a network of NLR proteins. Previous studies have shown that resistance to some pathogens requires alternative splicing [31,32], such as *RPS4* in *Arabidopsis* [33]; and splicing is used to truncate proteins like *RCT1* in *Medicago truncatula* [34]. Wheat has also shown evidence of alternative splicing of important resistance genes Lr10 and Sr35 [35,36]. Splicing patterns between wheat paralogs resulting from duplication also appear to be conserved. Stop codon-containing inter-exon regions can be included in the transcript to force a truncation of the protein. Truncated NB-ARCs may result in decoy proteins, where signaling function is lost but IDs ‘distract’ pathogenic effectors from functional target proteins. The potential involvement of alternative splicing is shown in Figure 4, integrated into the varied use of IDs in NLR proteins. In concurrence with our results, Yang et al. (2014) describe NLR alternative splicing as useful for regulating NLR autoinhibition or function in signal transduction, also detailing potential transcription factor activity [31]. Genes that have multiple copies of an ID can also regulate the number of copies in the protein through alternative splicing. Accession 7BL_AA1863630 encodes two transcripts, one with three WD domains (G-beta repeat) and the other with two, along with Coils and NB-ARCs. Accession 3B_AA0740740 encodes five transcripts that all contain two separate ribosome inactivating protein domains, four of which also have NB-ARCs. Alternative splicing may also allow the plant to select different localization for a gene product. For example, 4AL_AA0961460, encodes three transcripts, one with multiple transmembrane helices and a BED zinc finger ID, one lacking the transmembrane helices, and another lacking the transmembrane helices and the ID. The transmembrane-containing transcript is encoded by one and a half extra exons at the beginning of the transcript, with a total of five exons making up the gene.

**Figure 4.**
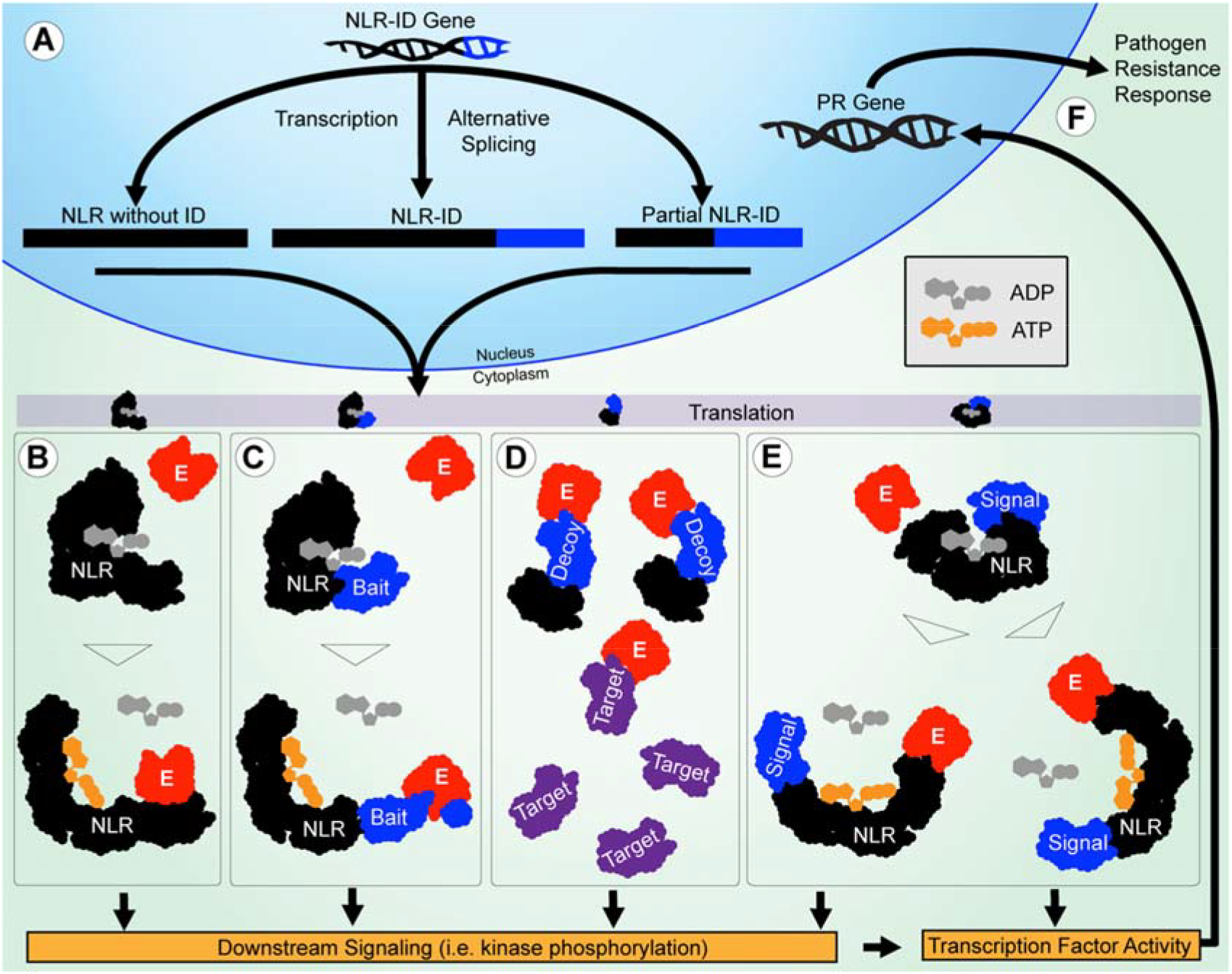
Potential roles of IDs in functional diversification of NLRs in pathogen resistance are illustrated. **(A)** The NLR-ID gene is alternatively spliced during transcription to include or exclude IDs. The NLR and ID sequences are shown in black and blue, respectively. **(B)** The NLRs without IDs function through effector-LRR binding to activate the protein, and trigger downstream signaling. Effectors are shown in red. **(C)** When IDs are used as baits, they mimic pathogen targets, and cause NLR activation after they are modified. **(D)** When IDs are used as decoys, they mimic pathogen targets to reduce effector interference in resistance signaling. The targets are shown in purple. **(E)** When IDs are used in signaling, they allow NLRs to act as signal transduction factors, less reliant on downstream signaling utilized by other NLRs. **(F)** Finally, transcription factor activity directly involving or triggered by NLRs causes PR genes to be expressed, leading to a resistance response.

Wheat expression data shows that there are differences in the expression of these alternative transcripts shown in Figure 3. We mined the expression values for the 54 transcripts present in Figure 3 from datasets present in the Wheat Gene Expression Atlas and NCBI databases. Expression data from Salcedo et al. (2017) shows that alternative splicing may result from different conditions [37]. At very least, these data provide support for Figure 3 accessions as actual alternative transcripts that can be measured experimentally. This data is present in Figure S1 and **Table S1**. In the Wheat Gene Expression Atlas data [38], several transcripts with different ID contents show differential expression in wheat tissues. For instance, both transcripts of 5AL_AA1195550 were expressed, much more for the WRKY-containing first transcript. Similar examples include 5AL_AA1200550 (kinase), 5BL_AA1321240 (kinase), 5BS_AA1380910 (LRR), 5DS_AA1479690 (kinase and coil), 7AS_AA1808020 (zinc finger), and 7DS_AA2051850 (B3). Accession 3B_AA0740740 shows evidence of higher expression of a truncated NLR-ID, lacking an NB-ARC. Several of these genes were more strongly expressed in the leaves, shoots, and spikes. One exception is 7DS_AA2051850, where the second B3-containing transcript is expressed much more in the roots than in any other tissue, and much higher than the other B3-lacking transcript in the roots. This gene may be involved in resistance to soil-borne pathogens. More data is required to conclusively show differential expression based upon certain treatments and conditions.

While wheat shows evidence of NLR-ID alternative splicing, barley may have evolved a more diverse set of transcripts for NLR-IDs. Several barley NLR-ID proteins have dozens of transcripts, with several of those allowing for alternative use of IDs in NLR proteins. Many barley genes have alternative transcripts that encode NLR-ID, just NLR, just ID, or lack both. For example, barley gene HORVU3Hr1G010980 encodes 16 transcripts, many of which contain NB-ARCs and two separate ribosome inactivating domains, one with just the NB-ARC, one just the ribosome inactivating domain, and one short transcript without any domains. Previous studies have identified barley *Mla* genes as utilizing alternative splicing for resistance [39]. Barley genes can also encode multiple IDs. HORVU3Hr1G037800 and HORVU7Hr1G000320 can encode transcripts containing NB-ARC domains, Kelch motifs, a Glutaredoxin domain, or a short transcript with none of those domains. HORVU1Hr1G079170 contains dozens of transcripts that can either encode NB-ARC and LRR protein or proteins with Glycoside hydrolase family 28 and TM helix proteins, potentially involved in two different regulatory pathways. HORVU7Hr1G120020 encodes 17 transcripts with NB-ARC, Coil, and Cleavage site for pathogenic type III effector avirulence factor Avr, with some transcripts only encoding the cleavage site with no NB-ARC, a possible decoy for pathogenic effectors. Barley gene HORVU5Hr1G085900 encodes 20 transcripts, some as NB-ARC-WRKY, just WSKY, and NB-ARC-WRKY-WSKY. Since some transcripts just contain the transcription factor domain, either this is functioning as a transcription factor, or it is a non-receptor decoy that reduces effector interference in WRKYs necessary for resistance response. This protein also contains a probable nuclear localization signal between the NB-ARC and the WRKY domains. Our data supports a previous prediction that alternative splicing may allow for differential cellular localization [31].

## 4. MATERIALS AND METHODS

*Triticum aestivum* protein sequences were downloaded using the Biomart application within the Ensembl Genomes [40] and Phytozome [41] databases. InterProScan annotations [42] were compiled and proteins containing NB-ARC domains (PF00931) were investigated. Pfam annotations not inherently part of NLR structure were assembled. Amino acid sequences and corresponding annotations were uploaded to the program Geneious [43] for sequence alignment, homology assessment, and motif visualization. The IDs were manually investigated to assess amino acid location, potential function, homology to proteins in other species, and presence in variant transcripts. Function was assessed partially through domain descriptions available through the Pfam database [44], allowing for inferences about domain activity. Genomes investigated for homology include: *Aegilops tauschii*, *Amborella trichopoda*, *Arabidopsis thaliana*, *Brachypodium distachyon*, *Hordeum vulgare*, *Musa acuminata*, *Oryza sativa*, *Setaria italica*, *Triticum urartu*, and *Zea mays*. Genomic data was not available for *Aegilops speltoides*, which is believed to be the contributor of wheat’s B genome. The Gene Structure Display Server 2.0 [45] was used to visualize alternative splicing of NLR-IDs. Wheat expression data was generated from datasets in NCBI and Wheat Gene Expression Atlas data [38].

## 5. CONCLUSIONS

We found that the diversity of integrated domains in NLRs corresponds directly to the multiple components utilized by plant cells to initiate resistance responses, such as kinases, transcription factors, hormone signaling receptors, and proteins involved in antimicrobial compound production. NLR-ID fusions give these immune receptors the potential to function as effector baits, decoys, and signal transduction factors. This functional diversification would allow plants to stop using intermediate signaling factors that effectors often target to inhibit resistance. Sequence homology both indicates that some IDs retain functionality and provides an explanation for ID origins as functional non-NLR proteins before their integration into NLRs. Using both genomic and expression data, we showed that plants likely utilize alternative splicing to regulate the inclusion or exclusion of IDs in NLR proteins and have compiled a list of experientially verified alternative transcripts that truncate or exclude these domains. This optional deployment of NLR-IDs constitutes an important defense strategy to deal with rapidly evolving pathogen effectors. Future studies should aim to characterize the structure of NLR-ID fusion proteins, demonstrate which IDs have retained enzymatic activity, and associate the expression of alternative transcripts with specific conditions.

## Supplementary Materials

Supplementary materials can be found at www.mdpi.com/xxx/s1.

**Figure S1.**
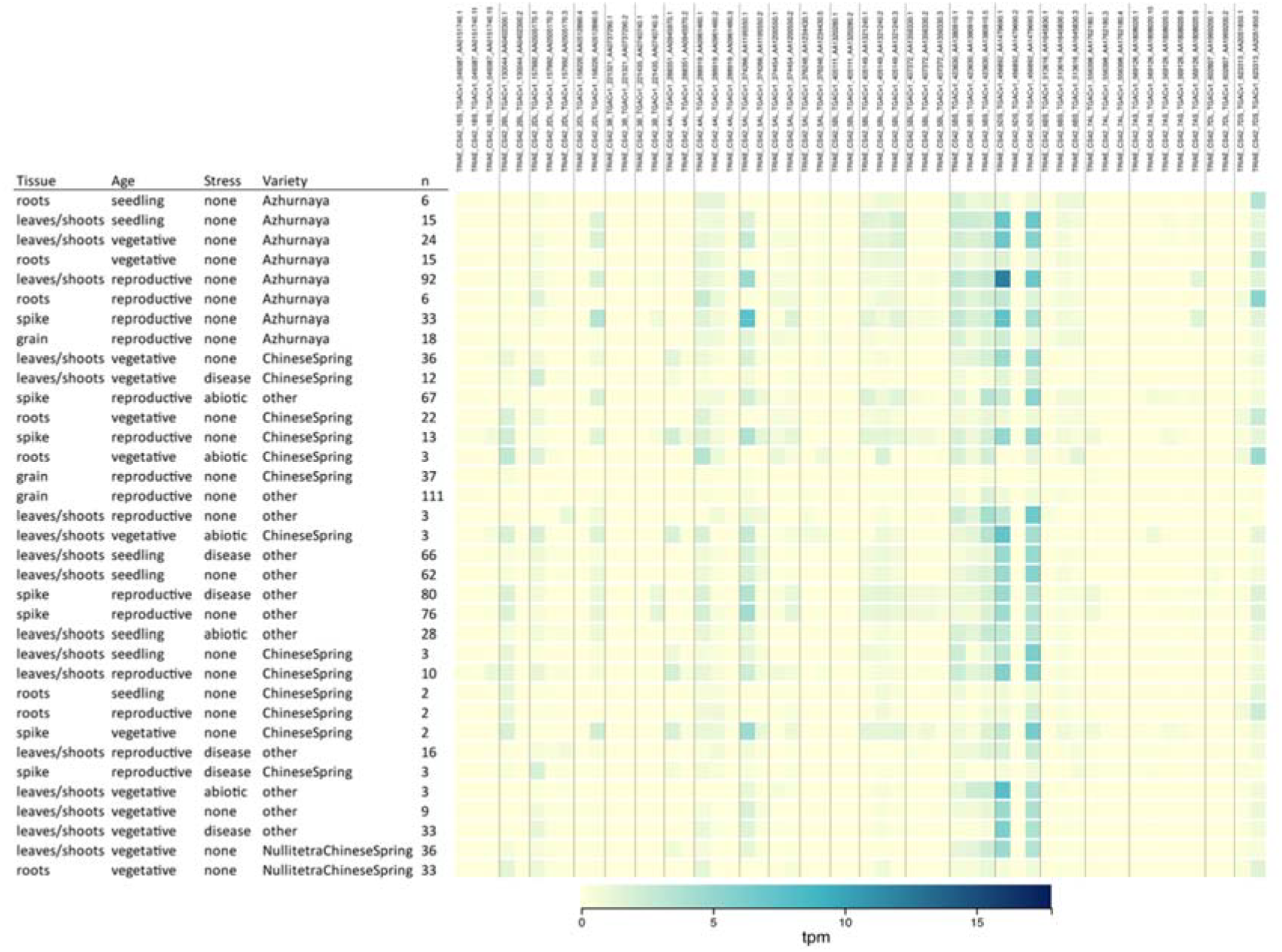
Expression of wheat NLR-ID alternative splicing candidate genes from Wheat Gene Expression Atlas. The 54 transcripts shown in Figure 3 were used to generate this heatmap visualization within the Wheat Gene Expression Atlas database. Visualization layout was made based upon expVIP within the database.

**Table S1.** Expression of wheat NLR-ID alternative splicing candidate genes available in Wheat Gene Expression Atlas and NCBI databases. The 54 transcripts shown in Figure 3 were acquired from the GSE106397 dataset in NCBI and the Wheat Gene Expression Atlas. Cells are colored by expression level, showing differential expression between alternative transcripts of the same gene.

[**Table S1 is now posted at** https://doi.org/10.6084/m9.figshare.8796449]

## AUTHOR CONTRIBUTIONS

EJA analyzed genomic data, interpreted results, and drafted the manuscript. MPN provided insight, constructive feedback, and rewrote some portions of the manuscript.

## ACKNOWLEDGMENTS

This project was supported by the South Dakota Agriculture Experiment Station (United States Department of Agriculture – National Institute of Food and Agriculture) hatch fund to MPN.

## REFERENCES

1. Jones, J.D.; Dangl, J.L. The plant immune system. Nature 2006, 444, 323–329.

2. Jones, J.D.; Vance, R.E.; Dangl, J.L. Intracellular innate immune surveillance devices in plants and animals. Science 2016, 354, aaf6395.

3. Shao, Z.Q.; Xue, J.Y.; Wu, P.; Zhang, Y.M.; Wu, Y.; Hang, Y.Y.; Wang, B.; Chen, J.Q. Large-scale analyses of angiosperm nucleotide-binding site-leucine-rich repeat genes reveal three anciently diverged classes with distinct evolutionary patterns. Plant Physiol 2016, 170, 2095–2109.

4. DeYoung, B.J.; Innes, R.W. Plant nbs-lrr proteins in pathogen sensing and host defense. Nature immunology 2006, 7, 1243.

5. Hayashi, N.; Inoue, H.; Kato, T.; Funao, T.; Shirota, M.; Shimizu, T.; Kanamori, H.; Yamane, H.; Hayano-Saito, Y.; Matsumoto, T. Durable panicle blast-resistance gene pb1 encodes an atypical cc-nbs-lrr protein and was generated by acquiring a promoter through local genome duplication. The Plant Journal 2010, 64, 498–510.

6. Kofoed, E.M.; Vance, R.E. Innate immune recognition of bacterial ligands by naips determines inflammasome specificity. Nature 2011, 477, 592.

7. Bonardi, V.; Cherkis, K.; Nishimura, M.T.; Dangl, J.L. A new eye on nlr proteins: Focused on clarity or diffused by complexity? Current opinion in immunology 2012, 24, 41–50.

8. Bonardi, V.; Dangl, J.L. How complex are intracellular immune receptor signaling complexes? Frontiers in plant science 2012, 3, 237.

9. Cesari, S.; Bernoux, M.; Moncuquet, P.; Kroj, T.; Dodds, P.N. A novel conserved mechanism for plant nlr protein pairs: The “integrated decoy” hypothesis. Frontiers in plant science 2014, 5.

10. Wu, C.-H.; Krasileva, K.V.; Banfield, M.J.; Terauchi, R.; Kamoun, S. The “sensor domains” of plant nlr proteins: More than decoys? Frontiers in plant science 2015, 6.

11. Sarris, P.F.; Cevik, V.; Dagdas, G.; Jones, J.D.; Krasileva, K.V. Comparative analysis of plant immune receptor architectures uncovers host proteins likely targeted by pathogens. BMC biology 2016, 14, 8.

12. Takken, F.L.; Goverse, A. How to build a pathogen detector: Structural basis of nb-lrr function. Current opinion in plant biology 2012, 15, 375–384.

13. Michelmore, R.W.; Christopoulou, M.; Caldwell, K.S. Impacts of resistance gene genetics, function, and evolution on a durable future. Annual review of phytopathology 2013, 51, 291–319.

14. Cui, H.; Tsuda, K.; Parker, J.E. Effector-triggered immunity: From pathogen perception to robust defense. Annual review of plant biology 2015, 66, 487–511.

15. Walker, J.E.; Saraste, M.; Runswick, M.J.; Gay, N.J. Distantly related sequences in the alpha-and beta-subunits of atp synthase, myosin, kinases and other atp-requiring enzymes and a common nucleotide binding fold. The EMBO journal 1982, 1, 945–951.

16. Gutterson, N.; Reuber, T.L. Regulation of disease resistance pathways by ap2/erf transcription factors. Current opinion in plant biology 2004, 7, 465–471.

17. Buscaill, P.; Rivas, S. Transcriptional control of plant defence responses. Current opinion in plant biology 2014, 20, 35–46.

18. Jonak, C.; Ökrész, L.; Bögre, L.; Hirt, H. Complexity, cross talk and integration of plant map kinase signalling. Current opinion in plant biology 2002, 5, 415–424.

19. Meyers, B.C.; Kozik, A.; Griego, A.; Kuang, H.; Michelmore, R.W. Genome-wide analysis of nbs-lrr– encoding genes in arabidopsis. The Plant Cell Online 2003, 15, 809–834.

20. Noutoshi, Y.; Ito, T.; Seki, M.; Nakashita, H.; Yoshida, S.; Marco, Y.; Shirasu, K.; Shinozaki, K. A single amino acid insertion in the wrky domain of the arabidopsis tir–nbs–lrr–wrky-type disease resistance protein slh1 (sensitive to low humidity 1) causes activation of defense responses and hypersensitive cell death. The Plant Journal 2005, 43, 873–888.

21. Lannoo, N.; Van Damme, E.J. Lectin domains at the frontiers of plant defense. 2014.

22. Xiang, Y.; Song, M.; Wei, Z.; Tong, J.; Zhang, L.; Xiao, L.; Ma, Z.; Wang, Y. A jacalin-related lectin-like gene in wheat is a component of the plant defence system. Journal of experimental botany 2011, 62, 5471–5483.

23. Esch, L.; Schaffrath, U. An update on jacalin-like lectins and their role in plant defense. International journal of molecular sciences 2017, 18, 1592.

24. Hwang, I.S.; Hwang, B.K. The pepper mannose-binding lectin gene cambl1 is required to regulate cell death and defense responses to microbial pathogens. Plant physiology 2011, 155, 447–463.

25. Sarris, P.F.; Duxbury, Z.; Huh, S.U.; Ma, Y.; Segonzac, C.; Sklenar, J.; Derbyshire, P.; Cevik, V.; Rallapalli, G.; Saucet, S.B. A plant immune receptor detects pathogen effectors that target wrky transcription factors. Cell 2015, 161, 1089–1100.

26. Le Roux, C.; Huet, G.; Jauneau, A.; Camborde, L.; Trémousaygue, D.; Kraut, A.; Zhou, B.; Levaillant, M.; Adachi, H.; Yoshioka, H. A receptor pair with an integrated decoy converts pathogen disabling of transcription factors to immunity. Cell 2015, 161, 1074–1088.

27. Date, S.V. The rosetta stone method. In Bioinformatics, Springer: 2008; pp 169–180.

28. Baggs, E.; Dagdas, G.; Krasileva, K. Nlr diversity, helpers and integrated domains: Making sense of the nlr identity. Current opinion in plant biology 2017, 38, 59–67.

29. Consortium, I.W.G.S. A chromosome-based draft sequence of the hexaploid bread wheat (triticum aestivum) genome. Science 2014, 345, 1251788.

30. Kroj, T.; Chanclud, E.; Michel-Romiti, C.; Grand, X.; Morel, J.B. Integration of decoy domains derived from protein targets of pathogen effectors into plant immune receptors is widespread. New Phytologist 2016, 210, 618–626.

31. Yang, S.; Tang, F.; Zhu, H. Alternative splicing in plant immunity. International journal of molecular sciences 2014, 15, 10424–10445.

32. Shang, X.; Cao, Y.; Ma, L. Alternative splicing in plant genes: A means of regulating the environmental fitness of plants. International journal of molecular sciences 2017, 18, 432.

33. Zhang, X.-C.; Gassmann, W. Rps4-mediated disease resistance requires the combined presence of rps4 transcripts with full-length and truncated open reading frames. The Plant Cell 2003, 15, 2333–2342.

34. Tang, F.; Yang, S.; Gao, M.; Zhu, H. Alternative splicing is required for rct1-mediated disease resistance in medicago truncatula. Plant molecular biology 2013, 82, 367–374.

35. Sela, H.; Spiridon, L.N.; Petrescu, A.J.; Akerman, M.; Mandel-gutfreund, Y.; Nevo, E.; Loutre, C.; Keller, B.; Schulman, A.H.; Fahima, T. Ancient diversity of splicing motifs and protein surfaces in the wild emmer wheat (triticum dicoccoides) lr10 coiled coil (cc) and leucine-rich repeat (lrr) domains. Molecular plant pathology 2012, 13, 276–287.

36. Saintenac, C.; Zhang, W.; Salcedo, A.; Rouse, M.N.; Trick, H.N.; Akhunov, E.; Dubcovsky, J. Identification of wheat gene sr35 that confers resistance to ug99 stem rust race group. Science 2013, 341, 783–786.

37. Salcedo, A.; Rutter, W.; Wang, S.; Akhunova, A.; Bolus, S.; Chao, S.; Anderson, N.; De Soto, M.F.; Rouse, M.; Szabo, L. Variation in the avrsr35 gene determines sr35 resistance against wheat stem rust race ug99. Science 2017, 358, 1604–1606.

38. Borrill, P.; Ramirez-Gonzalez, R.; Uauy, C. Expvip: A customisable rna-seq data analysis and visualisation platform opens up gene expression analysis. Plant Physiology 2016, pp. 01667.02015.

39. Halterman, D.A.; Wei, F.; Wise, R.P. Powdery mildew-induced mla mrnas are alternatively spliced and contain multiple upstream open reading frames. Plant physiology 2003, 131, 558–567.

40. Kersey, P.J.; Allen, J.E.; Armean, I.; Boddu, S.; Bolt, B.J.; Carvalho-Silva, D.; Christensen, M.; Davis, P.; Falin, L.J.; Grabmueller, C. Ensembl genomes 2016: More genomes, more complexity. Nucleic acids research 2015, 44, D574–D580.

41. Goodstein, D.M.; Shu, S.; Howson, R.; Neupane, R.; Hayes, R.D.; Fazo, J.; Mitros, T.; Dirks, W.; Hellsten, U.; Putnam, N. Phytozome: A comparative platform for green plant genomics. Nucleic acids research 2011, 40, D1178–D1186.

42. Jones, P.; Binns, D.; Chang, H.-Y.; Fraser, M.; Li, W.; McAnulla, C.; McWilliam, H.; Maslen, J.; Mitchell, A.; Nuka, G. Interproscan 5: Genome-scale protein function classification. Bioinformatics 2014, 30, 1236–1240.

43. Kearse, M.; Moir, R.; Wilson, A.; Stones-Havas, S.; Cheung, M.; Sturrock, S.; Buxton, S.; Cooper, A.; Markowitz, S.; Duran, C. Geneious basic: An integrated and extendable desktop software platform for the organization and analysis of sequence data. Bioinformatics 2012, 28, 1647–1649.

44. Finn, R.D.; Coggill, P.; Eberhardt, R.Y.; Eddy, S.R.; Mistry, J.; Mitchell, A.L.; Potter, S.C.; Punta, M.; Qureshi, M.; Sangrador-Vegas, A. The pfam protein families database: Towards a more sustainable future. Nucleic acids research 2015, 44, D279–D285.

45. Hu, B.; Jin, J.; Guo, A.-Y.; Zhang, H.; Luo, J.; Gao, G. Gsds 2.0: An upgraded gene feature visualization server. Bioinformatics 2014, 31, 1296–1297.

